# Performance benchmarking of GATK3.8 and GATK4

**DOI:** 10.1101/348565

**Authors:** Jacob R. Heldenbrand, Saurabh Baheti, Matthew A. Bockol, Travis M. Drucker, Steven N. Hart, Matthew E. Hudson, Ravishankar K. Iyer, Michael T. Kalmbach, Eric W. Klee, Eric D. Wieben, Mathieu Wiepert, Derek E. Wildman, Liudmila S. Mainzer

## Abstract

Use of the Genome Analysis Toolkit (GATK) continues to be the standard practice in genomic variant calling in both research and the clinic. Recently the toolkit has been rapidly evolving. Significant computational performance improvements have been introduced in GATK3.8 through collaboration with Intel in 2017. The first release of GATK4 in early 2018 revealed significant rewrites in the code base, as the stepping stone toward a Spark implementation. As the software continues to be a moving target for optimal deployment in highly productive environments, we present a detailed analysis of these improvements, to help the community stay abreast with changes in performance. We re-evaluated the options previously identified as advantageous, such as threading, parallel garbage collection, I/O options and data-level parallelization. Based on our results, we consider the performance and cost trade-offs of using GATK3.8 and GATK4 for different types of analyses.

## 1. Introduction

GATK3.8 is the latest release of the “traditional” Java-based GATK designed to work on regular servers or compute clusters. GATK4, first officially released in January of 2018, is meant to be eventually deployed on data analytics platforms. At present it contains both Spark and non-Spark implementations of many of the tools. Because most of the Spark tools were still in beta at the time of the initial release, we focused our testing on the non-Spark implementations. The two versions of GATK have some initial differences; for instance, the PrintReads tool from GATK3.8 has become ApplyBQSR in GATK4. When optimizing a workflow, one can perform two distinct optimizations, and we explore them both:

**maximizing speed:** minimize the time to process a single sample; useful in time-critical situations, i.e. when a patient has a critical or rapidly developing condition; see sections 2.2 through 2.6.

**maximizing throughput:** maximize the number of samples processed per unit time; cost-effective for routine analyses or large population studies; see section 2.7.

## 2. Methods and Results

### 2.1 Experimental setup

#### Software versions

GATK3.8 was downloaded from the Broad Institute’s software download page, build GATK-3.8-0-ge9d806836. Picard version 2.17.4 and GATK4.0.1.2 were downloaded from GitHub as pre-compiled jar files.

#### Tools

Our benchmarking focused on the GATK Best Practices [1,2] starting from the duplicate marking stage through variant calling. The MarkDuplicates tool is not part of GATK3 and was called from a separate toolkit, Picard. MarkDuplicates is included directly into GATK4. Realignment is no longer recommended, and was not tested. The base recalibration process consists of two tools, BaseRecalibrator and PrintReads/ApplyBQSR. The final tool we benchmarked was HaplotypeCaller, which is common to both versions of GATK.

#### Data

A dataset corresponding to whole genome sequencing (WGS) performed on NA12878 to ~20X depth was downloaded from Illumina BaseSpace on Dec 16, 2016. The paired-ended, 126 nt reads were aligned with BWA MEM [3] against the hg38 human reference (from the Oct 2017 GATK Bundle) and sorted with Novosort [4] prior to benchmarking. Some settings required multiple tests and measurements; in those cases we only used the reads that mapped to chromosome 21. For known sites, dbSNP build 146 was used.

#### Hardware

All tests were conducted on Skylake Xeon Gold 6148 processors with 40 cores, 2.40 GHz. Each node had 192 GB, 2666 MHz RAM. The nodes were stateless, connected to a network-attached IBM GPFS ver. 4.2.1 with custom metadata acceleration. The cluster used EDR InfiniBand with 100 Gb/sec bandwidth, 100 ns latency. Nodes ran Red Hat Enterprise Linux 6.9.

### 2.2 GATK3.8 Tool-level thread scalability

The non-Spark GATK4 version is entirely single-threaded, except for the PairHMM portion of HaplotypeCaller (section 2.5 below). Picard’s MarkDuplicates is also single-threaded. Thus, our thread scalability testing focused on the GATK3.8 tools. We measured the walltime for each tool when invoked with a certain thread count, in the range from 1 to 40. HaplotypeCaller has two types of threads: nt and nct. We kept nt at 1 and modified nct. When one thread is reported for HaplotypeCaller, one thread of each type was used.

The tools respond differently to multithreading. Both BaseRecalibrator and HaplotypeCaller experience a 5-fold speedup compared to a single-threaded run when using 16 threads, but do not scale beyond that (Figure 1a). PrintReads gains an initial improvement with 3 threads (the apparent optimum for our dataset), and experiences degraded performance at higher thread counts (Figure 1b).

**Figure 1.**
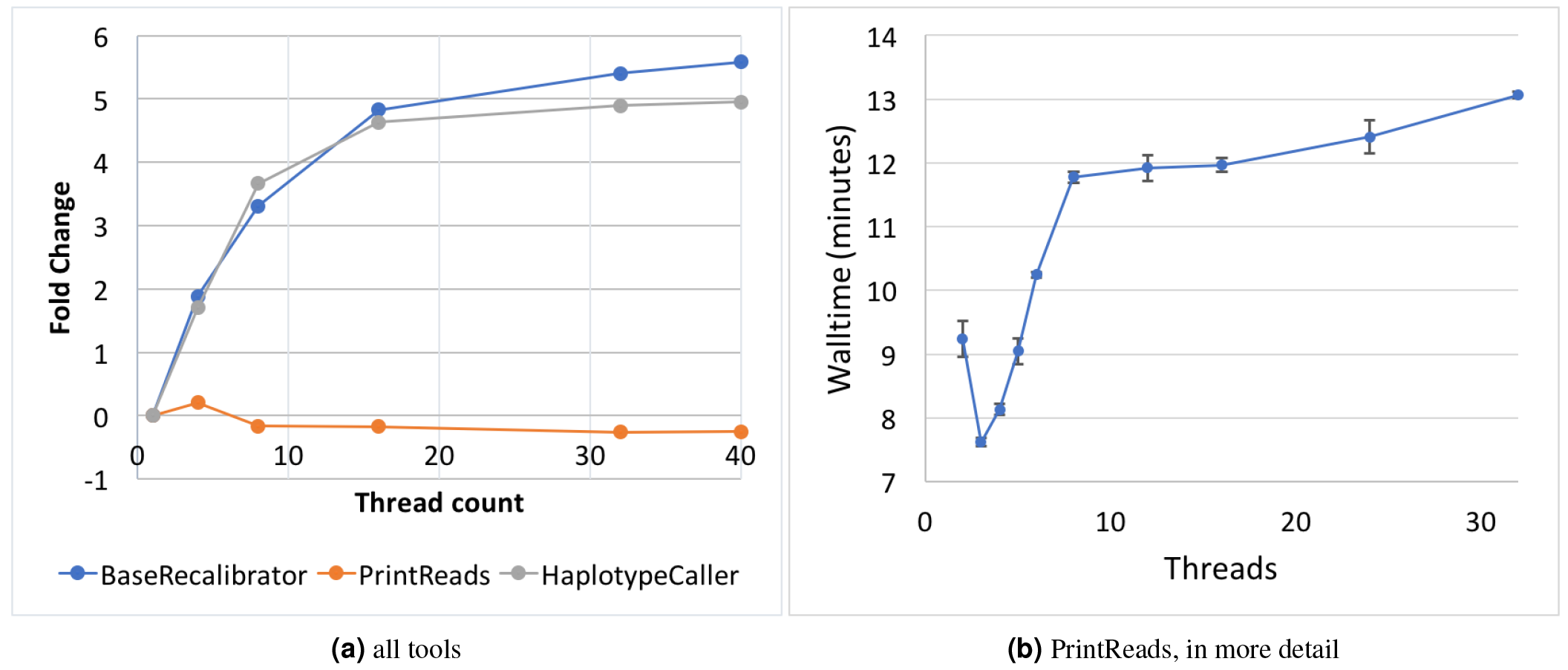
GATK3.8 Thread Scalability. (a) Sample: NA12878 WGS. Fold change refers to the fold difference in walltime between the new measurement when compared to the performance with a single thread ((*newtime ‒ baselinetime*)/*baselinetime*). (b) Sample: NA12878 chr 21. Error bars denote 1 SD around the mean of three replicates.

### 2.3 GATK4 Parallel garbage collection

A previous study [5] found that enabling Java parallel garbage collector (PGC) with up to 32 threads improved the walltime of GATK3.7. We explored this effect in the GATK4 tools.

The flags enabling PGC are passed to the GATK4 launch script via the “‒java-options” flag:

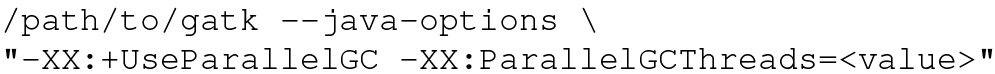

We found that enabling PGC for either ApplyBQSR or HaplotypeCaller had no impact or even degraded performance, depending on the number of threads used (data not shown). However, in MarkDuplicates using 2-4 PGC threads provided optimal performance (Figure 2a). For BaseRecalibrator, there is much more variability that we could not link to the state of the cluster (Figure 2b). The optimal thread choice appears to be around 24 threads, but the high walltimes at thread counts close to 24 suggest that it may be more reliable to 1) perform a similar thread count sweep on one’s own system to find the optimum, or 2) leave parallel garbage collection off to avoid one of the sub-optimal thread counts.

**Figure 2.**
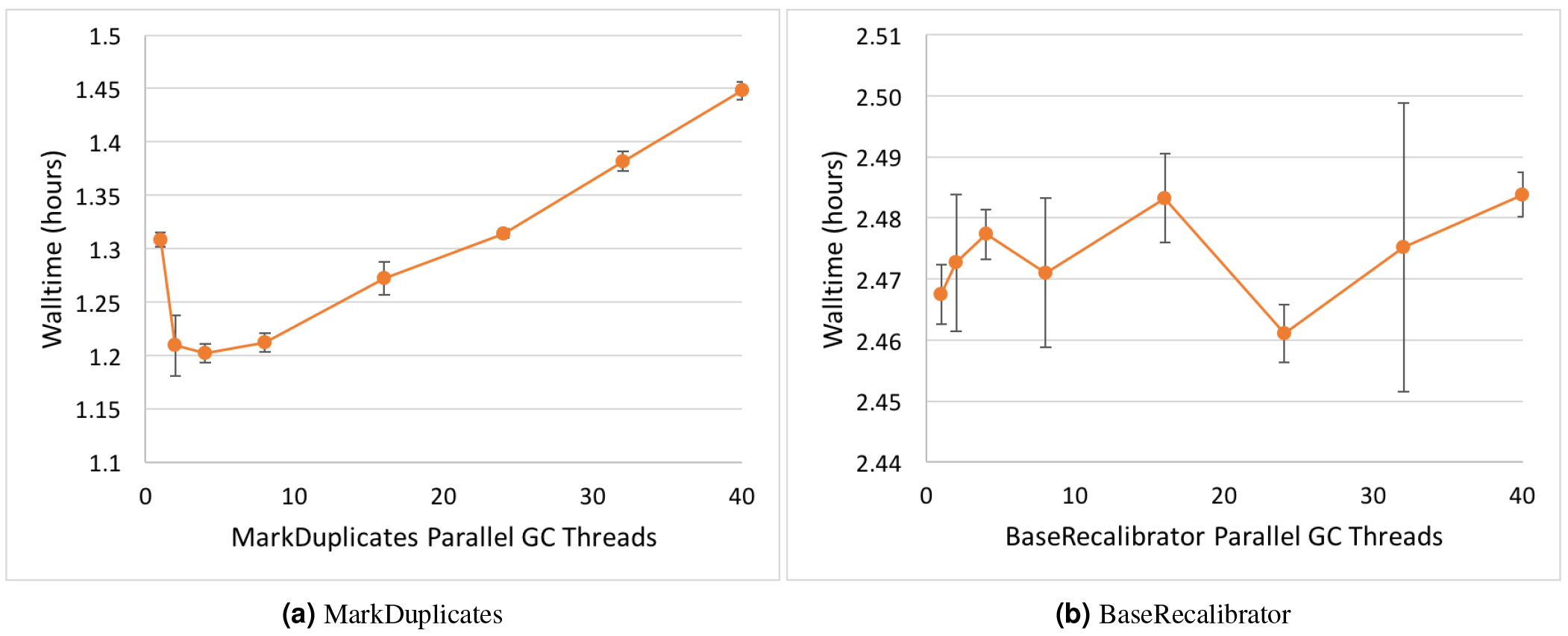
GATK4 thread scalability for Java parallel garbage collection. Sample: NA12878 WGS. The measurements at 1 PGC thread represent the default, meaning that PGC is not enabled. Error bars denote SD around the mean of three replicates.

We took a cursory look at PGC scalability in GATK3.8 and did not find significant improvements. In Picard’s MarkDuplicates, the optimum lies at approximately 2 PGC threads.

### 2.4 Asynchronous I/O in GATK 4

GATK4 has two types of asynchronous read/write options: Samtools I/O and Tribble I/O. “Tribble” is a specialized data format, mainly used for index files. To enable async I/O, one must edit the following variables in a gatk-properties file, located at `src/main/resources/org/broadinstitute/hellbender/utils/config/GATKConfig.properties` in the GATK GitHub repository:

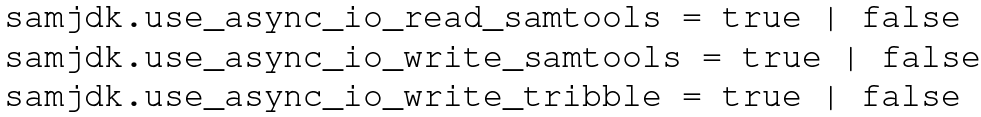

The properties file is passed to GATK with the ‘‒gatk-config-file’ flag. Because GATK4 MarkDuplicates is just a port of Picard’s tool of the same name, it does not accept a configuration file. We ran HaplotypeCaller with a single thread for this series of tests.

We found it best to enable asynchronous I/O for Samtools reading and writing and disable it for Tribble I/O (Table 1).

**Table 1.** Effects of asynchronous I/O settings on walltime (hours) in GATK4. Sample: NA12878 WGS

| Tool Name | Async I/O activated? |  |  |
| --- | --- | --- | --- |
|  | no | all | only for samtools I/O |
| BaseRecalibrator | 4.07 | 2.95 | 2.88 |
| ApplyBQSR | 2.38 | 2.07 | 2.08 |
| HaplotypeCaller | 17.25 | 17.31 | 17.08 |

### 2.5 PairHMM Scalability in GATK4 HaplotypeCaller

Intel partnered up with the Broad Institute to create the Genomics Kernel Library (GKL), which includes key optimizations to the HaplotypeCaller algorithm. The library introduces AVX optimized versions of the PairHMM and Smith-Waterman algorithms. Additionally, OpenMP support was added to the PairHMM algorithm to enable multithreading. While the library was developed to be used in GATK4, the AVX capabilities were back propagated to GATK3.8 as well.

The pre-built GATK4 that we downloaded from the repository was already configured to automatically detect hardware support for AVX. On our Skylake architecture, AVX-512 was utilized automatically.

The multi-threaded implementation of the PairHMM algorithm can be enabled with the following flags:

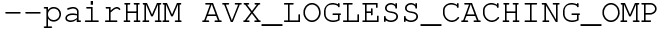

and

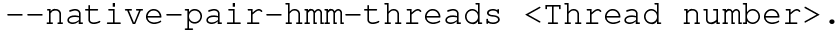

The optimum for GATK4 HaplotypeCaller seems to be around 10 threads (Figure 3).

**Figure 3.**
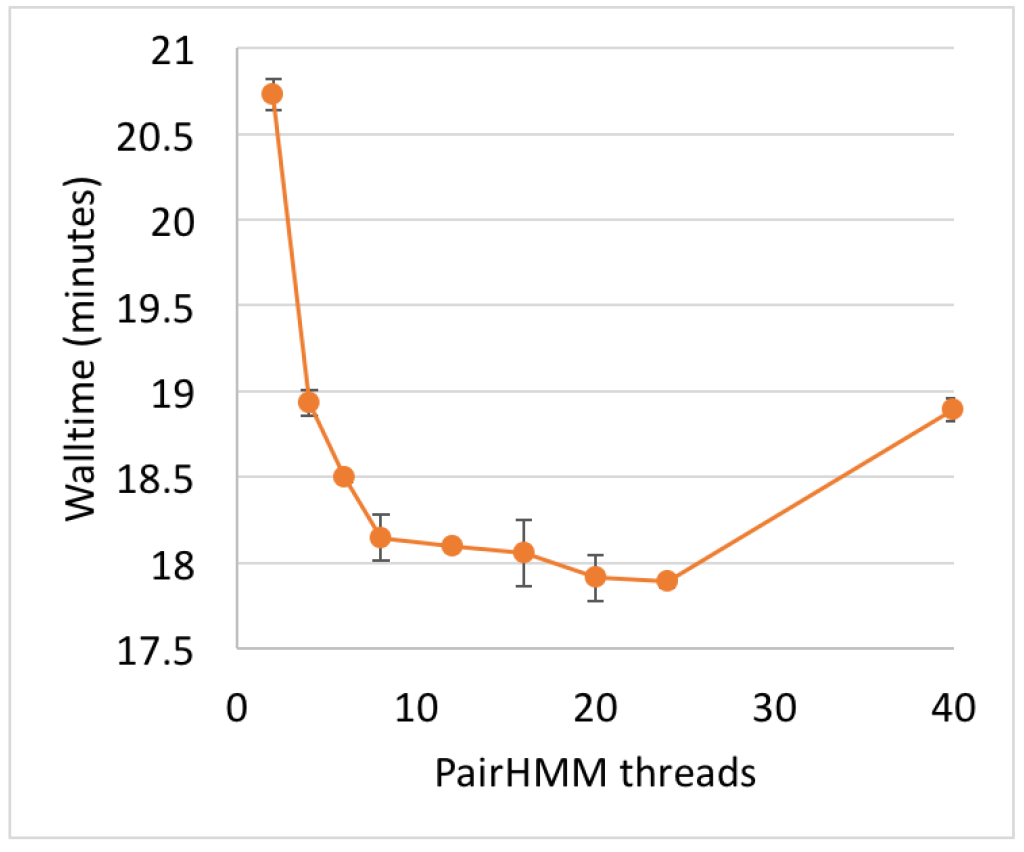
GATK4 thread scalability in HaplotypeCaller. Sample: NA12878 chr21. Error bars denote 1 SD around the mean of three replicates.

### 2.6 Splitting by chromosome

To achieve the greatest speedup, it is often efficient to split data by chromosome and process each interval in parallel. Here, we split the aligned sorted BAM into varying numbers of roughly equal-size chunks (Table 2) by using the GATK interval flag (-L) to observe how splitting affected walltime. The chunks were either kept on the same node for maximal utilization of cores (“within-node” parallelization) or spilled to more nodes for even shorter walltime (“across-node” parallelization).

**Table 2.**
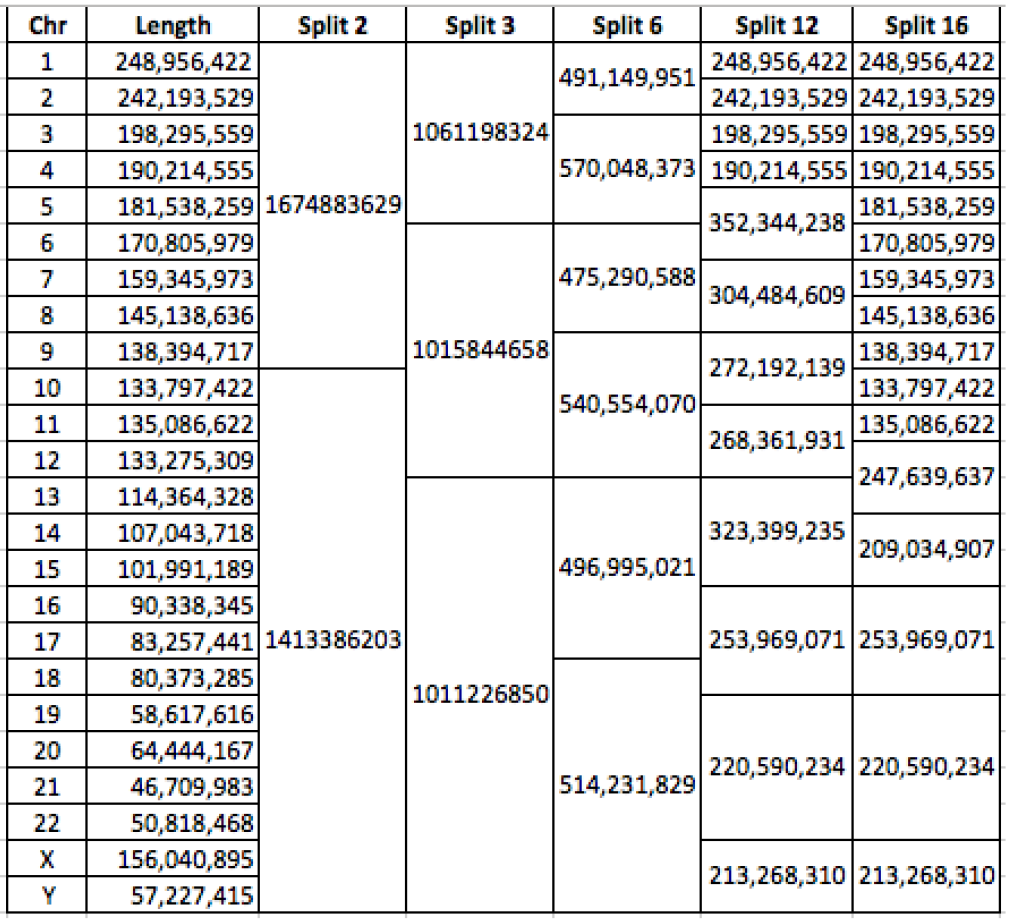
Chunking of the genome by chromosomes.

The previously discussed optimizations were applied in these experiments for both GATK3.8 and GATK4. For “within-node splitting,” we strove to optimally fill up our 40-core Skylake nodes by adjusting optimization parameters based on the number of chunks being processed in parallel within the node. For example, in GATK3.8 the optimal thread count for a tool may be around 10 threads, but we set the thread count for each chunk to 3 when the input is split into 12 chunks, while keeping all computations on the same node. Parallel garbage collection degrades the performance of BaseRecalibrator at lower thread counts and was therefore not used in the splitting experiments. Parallel GC was used with MarkDuplicates, but with only 2 threads, as that was optimal.

#### GATK3.8 results

For within-node parallelization beyond three chunks, the benefit of splitting the data begins to be counteracted by the degradation in performance caused by decreasing the thread count of each tool (Figure 4a). Thus it makes sense to spread execution over multiple nodes. We tested processing 6 chunks on 2 nodes, and 12 chunks on 4 nodes - thus keeping to 3 chunks per node (Figure 4b). This further reduced the total walltime, although perhaps at a higher compute cost.

**Figure 4.**
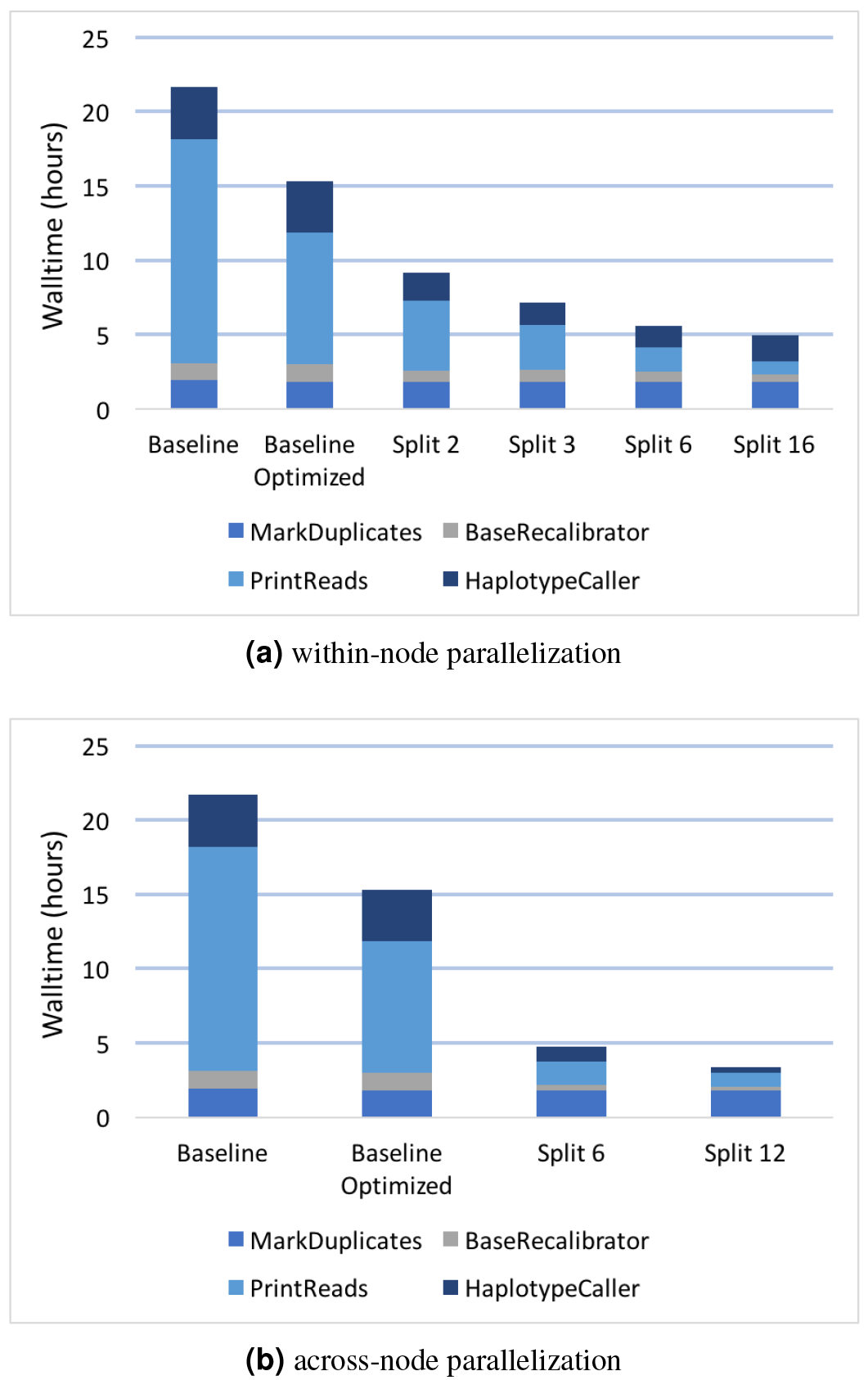
Effects of data-level parallelization in GATK3.8. Sample: NA12878 WGS. The “Baseline” was a naive approach where we gave each tool 40 threads (1 thread per core). The “Baseline Optimized” gave each tool 40 threads, except for PrintReads, which utilized 3. MarkDuplicates and BaseRecalibrator were given 2 and 20 parallel garbage collection threads, respectively. “Split 2,” “Split 3,” etc. means that the aligned sorted BAM was split into 2, 3, etc. chunks, as shown in Table 2. Panel (a) shows experiments with chunks computing on the same node. In panel (b) computation was spread across nodes in groups of 3 chunks per node.

#### GATK4 results

Splitting the aligned sorted BAM into chunks is simple in GATK4, as the only multithreaded tool is HaplotypeCaller. We again split into 2, 3, 6, and 16 chunks, which were kept on the same node, and the PairHMM thread count for HaplotypeCaller was adjusted accordingly (Figure 5). In contrast to the results we observed for GATK3.8, the walltime keeps improving when splitting all the way down to 16 chunks.

**Figure 5.**
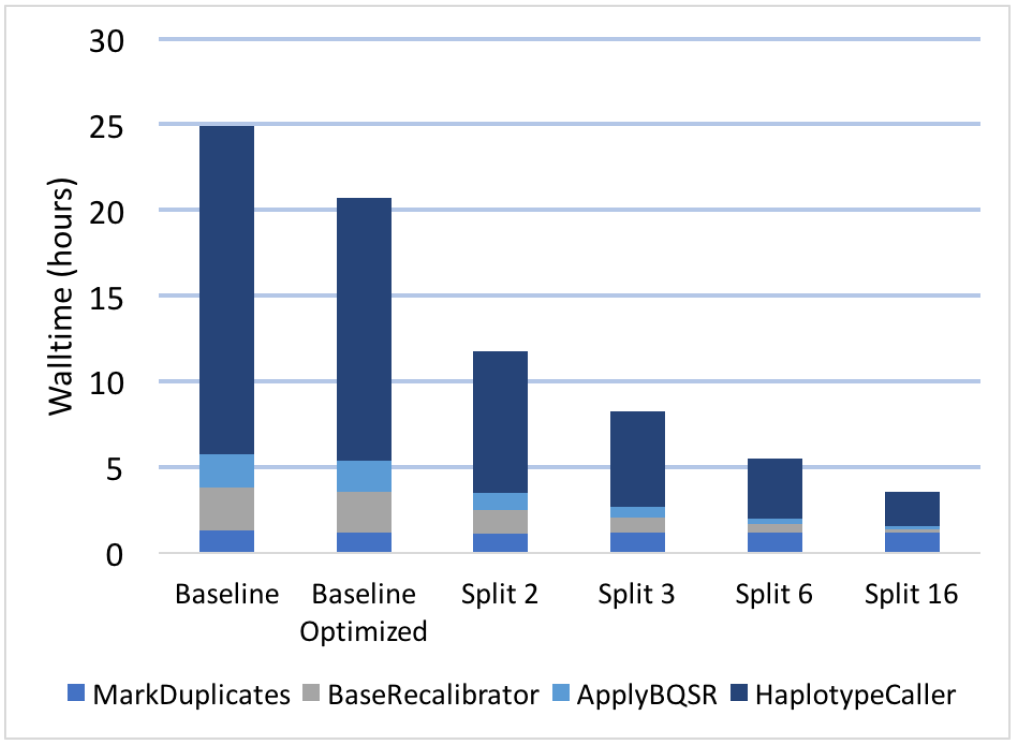
Effects of data-level parallelization in GATK 4. All compute was kept within the same node. Sample: NA12878 WGS. “Split 2,” “Split 3,” etc. means that the aligned sorted BAM was split into 2, 3, etc. chunks, as shown in Table 2.

### 2.7 Throughput

When optimizing throughput, one is maximizing the number of samples processed per unit time, albeit at the cost of higher walltime per sample. Because GATK4 is at present single-threaded by design, it lends itself extremely well to this kind of optimization. We created 40 copies of the NA12878 aligned sorted BAM file and processed them in parallel on a single 40-core node (Figure 6). The overall walltime does increase as one adds more samples to a node, probably due to contention for memory access and possibly disk I/O. However, the overall throughput increases substantially up until around 20 samples per node. Placing more than 20 samples on a 40-core Skylake node is probably not cost-effective.

**Figure 6.**
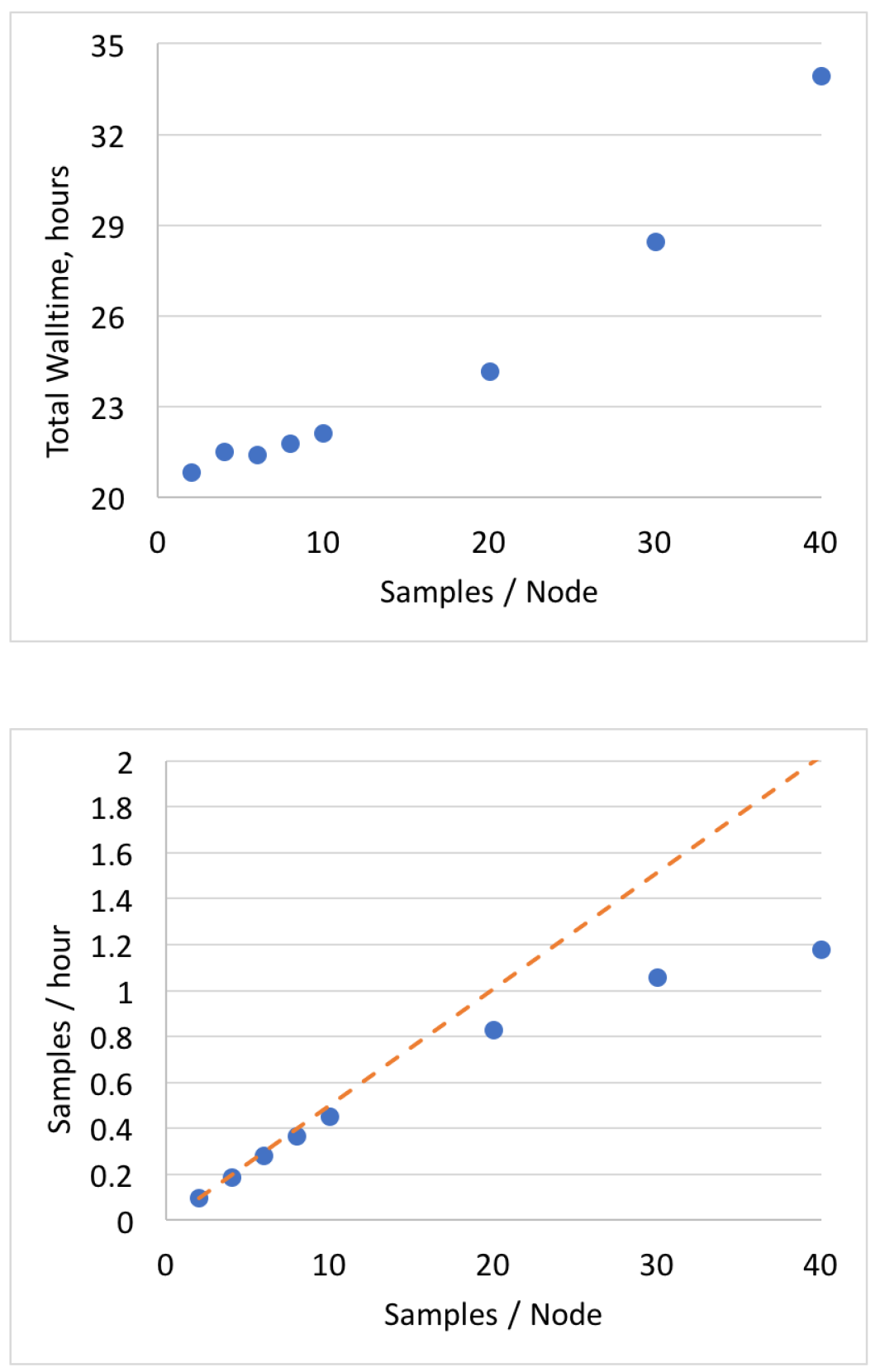
GATK4 throughput testing, measuring total walltime when running multiple samples simultaneously on the same node. Sample: NA12878 WGS.

## 3. Summary and Discussion

The tested optimizations intended to speed up computation in individual GATK tools are summarized in Table 3. When applied together, these optimizations significantly reduce the walltime on NA12878 WGS 20X (no splitting by chromosome). In GATK3.8 the MarkDuplicates → BaseRecalibrator → PrintReads → HaplotypeCaller walltime went from 21.7 hours down to 15.3 hours (29.3% improvement). In GATK4 the MarkDuplicates → BaseRecalibrator → ApplyBQSR → HaplotypeCaller walltime went from 24.9 hours to 20.7 hours (16.9% improvement). Note that the walltime is fairly comparable between the two GATK versions despite the single-threaded nature of GATK4, highlighting the performance optimizations introduced into that new release due to complete rewrite of many portions of the code.

**Table 3.**
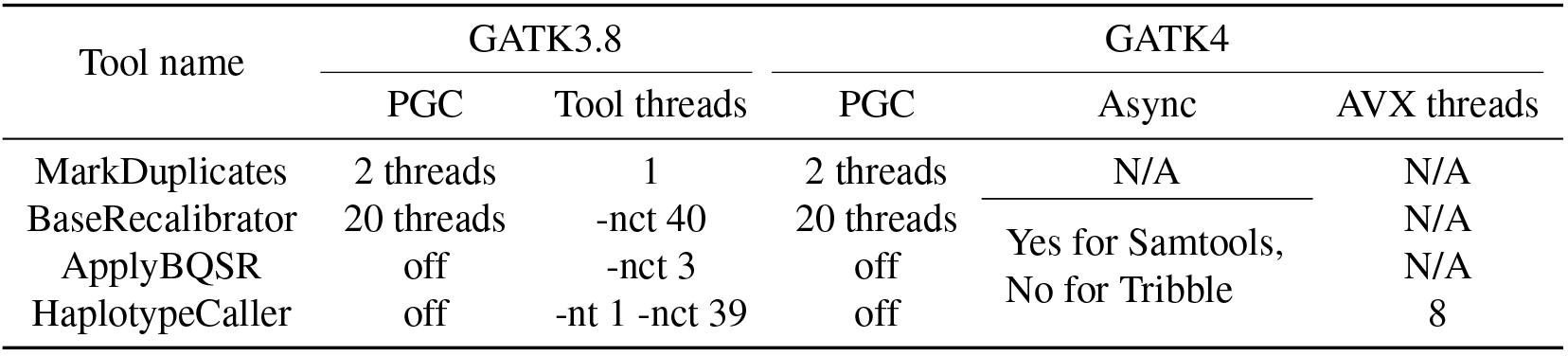
Summary of optimized parameter values.

Further walltime improvement can be achieved via splitting the aligned sorted BAM by chromosome. In GATK3.8 the walltime is reduced down to 5 hours when BAM is split into 16 chunks running on the same node ‒ a 76.9% improvement relative to the unoptimized, unsplit configuration. Further benefit can be achieved by splitting into 12 chunks across 4 nodes: down to 3.4 hours (84.3% total improvement). A similar walltime of 3.6 hours is accomplished in GATK4 by splitting into 16 chunks running on the same node ‒ potentially a very cost-effective solution.

To assess the financial costs and benefits resulting from the various configurations of the pipeline, we calculated the dollar amount for our runs based on AWS pricing. All our nodes are built with 40-core Skylake CPUs and 192 GB of RAM. This does not exactly match any of the AWS Skylake instances: c5.9xlarge gives 36 cores and 72 GB of RAM, and c5.18xlarge gives 72 cores and 144 GB of RAM. Our optimizations do aim to maximally pack our nodes with processes, but 72 GB of RAM would probably be insufficient for some high-throughput configurations. Thus Table 4 gives cost estimates for both types of instances, with the understanding that true values are somewhere in between.

**Table 4.** Financial costs per sample when running an optimized pipeline, based on AWS on-demand pricing as of June 2018: c5.9xlarge at $1.53 per hour and c5.18xlarge at $3.06 per hour. Configurations are sorted by cost.

| GATK version | Splitting | Samples | Nodes | Walltime, hrs | c5.9xlarge | c5.18xlarge |
| --- | --- | --- | --- | --- | --- | --- |
| GATK 4.0.1.2 | no splitting | 1 | 1 | 20.7 | \$31.7 | \$63.3 |
| GATK 3.8 | no splitting | 1 | 1 | 15.3 | \$23.4 | \$46.8 |
| GATK 3.8 | 12 chunks | 1 | 4 | 3.4 | \$20.8 | \$41.6 |
| GATK 3.8 | 6 chunks | 1 | 2 | 4.7 | \$14.4 | \$28.8 |
| GATK 3.8 | 16 chunks | 1 | 1 | 5.0 | \$7.7 | \$15.3 |
| GATK 4.0.1.2 | 16 chunks | 1 | 1 | 3.6 | \$5.5 | \$11.0 |
| GATK 4.0.1.2 | no splitting | 40 | 1 | 34.1 | \$1.3 | \$2.6 |

The data emphasize the trade-off between speed and per-sample cost of the analysis. To achieve the two types of optimizations outlined in the Introduction:

**maximizing speed:** to minimize the time to process a single sample, useful in time-critical situations, i.e. when a patient has a critical or rapidly developing condition, use GATK3.8 by splitting the sample into 12 chunks and computing across 4 nodes; resultant walltime is 3.4 hours at the cost of $41.60 on c5.18xlarge.
**maximizing throughput:** to maximize the number of samples processed per unit time, cost-effective for routine analyses or large population studies, use GATK4.0.1.2 by running 40 samples on one node; total walltime is 34.1 hours with 1.18 samples processed per hour at the cost of $2.60 per sample.

## Acknowledgments

This work was a product of the Mayo Clinic and Illinois Strategic Alliance for Technology-Based Healthcare. Major funding was provided by the Mayo Clinic Center for Individualized Medicine and the Todd and Karen Wanek Program for Hypoplastic Left Heart Syndrome. We thank the Interdisciplinary Health Sciences Institute, UIUC Institute for Genomic Biology and the National Center for Supercomputing Applications for their generous support and access to resources. We particularly acknowledge the support of Keith Stewart, M.B., Ch.B., Mayo Clinic/Illinois Grand Challenge Sponsor and Director of the Mayo Clinic Center for Individualized Medicine. Many thanks to the GATK team at the Broad Institute for their consultation and advice on the internals of GATK. Special gratitute to Katherine Kendig and Amy Weckle for managing the project and proofreading this text.

